# Similar neuronal imprint and absence of cross-seeded partner fibrils in α-synuclein aggregates from MSA and Parkinson’s disease brains

**DOI:** 10.1101/2021.06.22.449410

**Authors:** Florent Laferrière, Stéphane Claverol, Erwan Bezard, Francesca De Giorgi, François Ichas

**Affiliations:** CNRS, Institut des Maladies Neurodégénératives, UMR 5293, Bordeaux, France; Université de Bordeaux, Institut des Maladies Neurodégénératives, UMR 5293, Bordeaux, France; Plateforme Proteome, Univ. Bordeaux, Bordeaux, France; INSERM, Laboratoire de Neurosciences Expérimentales et Cliniques, U-1084, Université de Poitiers, Poitiers, France

**Author notes:** Co-senior authorship (Francesca De Giorgi, François Ichas). Correspondence to: - Dr. Florent Laferrière, Institut des Maladies Neurodégénératives, Centre Broca, Nouvelle-Aquitaine, Université de Bordeaux, 146 rue Léo Saignat 33000 Bordeaux, France.,., - Dr. François Ichas, Institut des Maladies Neurodégénératives, Centre Broca Nouvelle-Aquitaine, Université de Bordeaux, 146 rue Léo Saignat 33000 Bordeaux, France.,.

**Keywords:** alpha-synuclein, synucleinopathy, Lewy bodies, glial cytoplasmic inclusions, cross-seeding

## Abstract

Aggregated alpha-synuclein (α-syn) is a principal constituent of Lewy bodies (LBs) and glial cytoplasmic inclusions (GCIs) observed respectively inside neurons in Parkinson’s disease (PD) and oligodendrocytes in multiple system atrophy (MSA). Yet, the cellular origin, the pathophysiological role, and the mechanism of formation of these inclusions bodies (IBs) remain to be elucidated. It has recently been proposed that α-syn IBs eventually cause the demise of the host cell by virtue of the cumulative sequestration of partner proteins and organelles. In particular, the hypothesis of a local cross-seeding of other fibrillization-prone proteins like tau or TDP-43 has also been put forward. We submitted sarkosyl-insoluble extracts of post-mortem brain tissue from PD, MSA and control subjects to a comparative proteomic analysis to address these points. Our studies indicate that: i) α-syn is by far the most enriched protein in PD and MSA extracts compared to controls; ii) PD and MSA extracts share a striking overlap of their sarkosyl-insoluble proteomes, consisting of a vast majority of mitochondrial and neuronal synaptic proteins, and (iii) other fibrillization-prone protein candidates possibly cross-seeded by α-syn are neither found in PD nor MSA extracts. Thus, our results (i) support the idea that pre-assembled building blocks originating in neurons serve to the formation of GCIs in MSA, (ii) show no sign of amyloid cross-seeding in either synucleinopathy, and (iii) point to the sequestration of mitochondria and of neuronal synaptic components in both LBs and GCIs.

## INTRODUCTION

Synucleinopathies, such as Parkinson’s disease (PD), dementia with Lewy Bodies (DLB), or multiple system atrophy (MSA), are a group of neurodegenerative disorders of unknown etiology with very diverse clinical and pathological presentations. However, these diseases share certain standard features, the most significant being the aggregation of alpha-synuclein (α-syn), present in cytoplasmic inclusions bodies (IBs) in the cells of affected brain regions. These IBs are divergent in nature and cellular localization across these synucleinopathies. Lewy bodies (LBs) are found in neurons for PD and DLB, while glial cytoplasmic inclusions (GCIs) are detected in oligodendrocytes for MSA^1,2^. LBs are eosinophilic inclusions comprising filamentous structures in a dense core surrounded by a peripheral halo^3^ containing numerous membranous components and dysmorphic organelles^4^. GCIs are packed triangle or sickle-shaped, filamentous structures^5^. α-syn is the principal filament constituent of both IBs^2,5,6^.

Besides their histopathological characterization, the origins, the mechanisms of formation, and the pathophysiological role of IBs remain unelucidated. In agreement with early studies showing that IB-harboring neurons show no apparent signs of apoptosis^7,8^ and that there is no correlation between the extent of LB pathology and the extent of neuronal depletion in PD^9–11^, it appears likely that the α-syn fibrils accumulated in the IBs are not toxic by themselves, and that their compaction represents the result of a neutralization process rather. Instead, the concomitant and cumulative incorporation of mitochondria and proteins essential for cell function into the growing IBs could sporadically be detrimental to the host cell^12^. At the same time, amyloidogenic partner proteins like tau or TDP-43 could also, in some instances, be locally cross-seeded and cause the cell’s demise^13–15^. In both cases, the catabolism of IBs released by the dying host cells in the extracellular space could then cause a local leak of small IB fragments and α-syn fibrils from the IB mass, fragments that in turn would be capable of being taken up by the neighboring cells in which they would trigger a *de novo* aggregation of endogenous α-syn, the formation of novel IBs, and so on. Indeed, in mice, the intracerebral injection of IBs extracted from human DLB brains leads to the development and to the spread of a synucleinopathy with neuropathological and cytological features that are strikingly identical to the one observed after injections of pure recombinant α-syn fibrils^16,17^. Thus, if this IB-dependent intercellular spread mechanism holds, it is tempting to speculate that the composition of IBs present in a host cell could reflect the history of the spread, i.e., show proteins of the donor cell transferred together with the IB fragment seeds and eventually associated with the α-syn fibrils mass in the neo-formed IB. Such a theoretical possibility of retracing the spread history of IBs could probably help to solve the enigma concerning GCIs in MSA. These inclusions feature a high load of fibrillar α-syn, which contrasts with the very low physiological abundance of the protein in mature oligodendrocytes^18,19^. To date, the origin of these oligodendroglial IBs remains to be unraveled.

Proteomic analysis of fractions enriched in IBs from PD and MSA brain samples by different methods identified hundreds of proteins besides α-syn^3,5,6,20–27^. Their extraction was achieved by various methods such as laser capture microdissection^24^, density sucrose or Percoll gradients^27^, followed by FACS-sorting^21^ or immunocapture on magnetic beads^20,22,26^, with optional partial proteolytic digestion^6,25^. The extracted proteins include structural and cytoskeletal elements (neurofilaments, tubulins, TPPP/p25), α-synuclein-binding proteins (14-3-3, synphillin-1), ubiquitin-proteasome system components, α-ß-crystallin, heat shock proteins, or DJ-1. Recently, the list was refined and demonstrated a high burden of synaptic vesicle-related proteins within the IBs^25^. However, these candidates’ relevance in the disease pathogenesis is limited by the experimental procedures’ ability to isolate IBs from the surrounding subcellular structures and proteins.

To this end, we used a different approach in the present study. We purified and isolated aggregated α-syn and its associated insoluble proteomes from PD and MSA brains using their sarkosyl-insolubility. We used Sarkospin, a previously developed procedure for purifying pathological protein aggregates by sedimentation^17,28,29^. By adapting Sarkospin to α-syn, we sought to specifically isolate the aggregated forms of the protein from their physiological monomeric and oligomeric counterparts. The latter separation allowed us to scrutinize the accompanying insoluble proteomes associated with each disease and shed light on the cellular origin and the components of IBs.

## RESULTS

### Extraction of pathological alpha-synuclein from synucleinopathy brains and separation from its regular counterparts by an adapted Sarkospin procedure

Intending to extract and identify the insoluble proteomes associated with aggregated α-syn for each synucleinopathy, we adapted the Sarkospin procedure to the purification of α-syn pathological assemblies found in human post-mortem tissue samples of PD and MSA subjects. We previously developed this method to extract aggregated TDP-43^29^. It was adapted by using ultracentrifugation of the sample mixed within a sucrose cushion after its solubilization in sarkosyl at 37°C with simultaneous nuclease treatment (Fig. 1a). The resulting supernatant contains sarkosyl-soluble material. The insoluble entities are collected in a dry pellet that is compatible with a further identification or quantification of its protein content by immunoblotting or mass spectrometry. Above all, the extraction of pathological assemblies through insolubility allows a direct comparison to control samples by applying the same procedure to healthy brain samples. We thus proceeded with parallel Sarkospin extractions, sedimentations, and filter-trap assays of sarkosyl-insoluble aggregates (Fig. 1b, c) from the brains of human control, PD, and MSA subjects (n=3 independent subject brains per control or disease group).

**Fig. 1.**
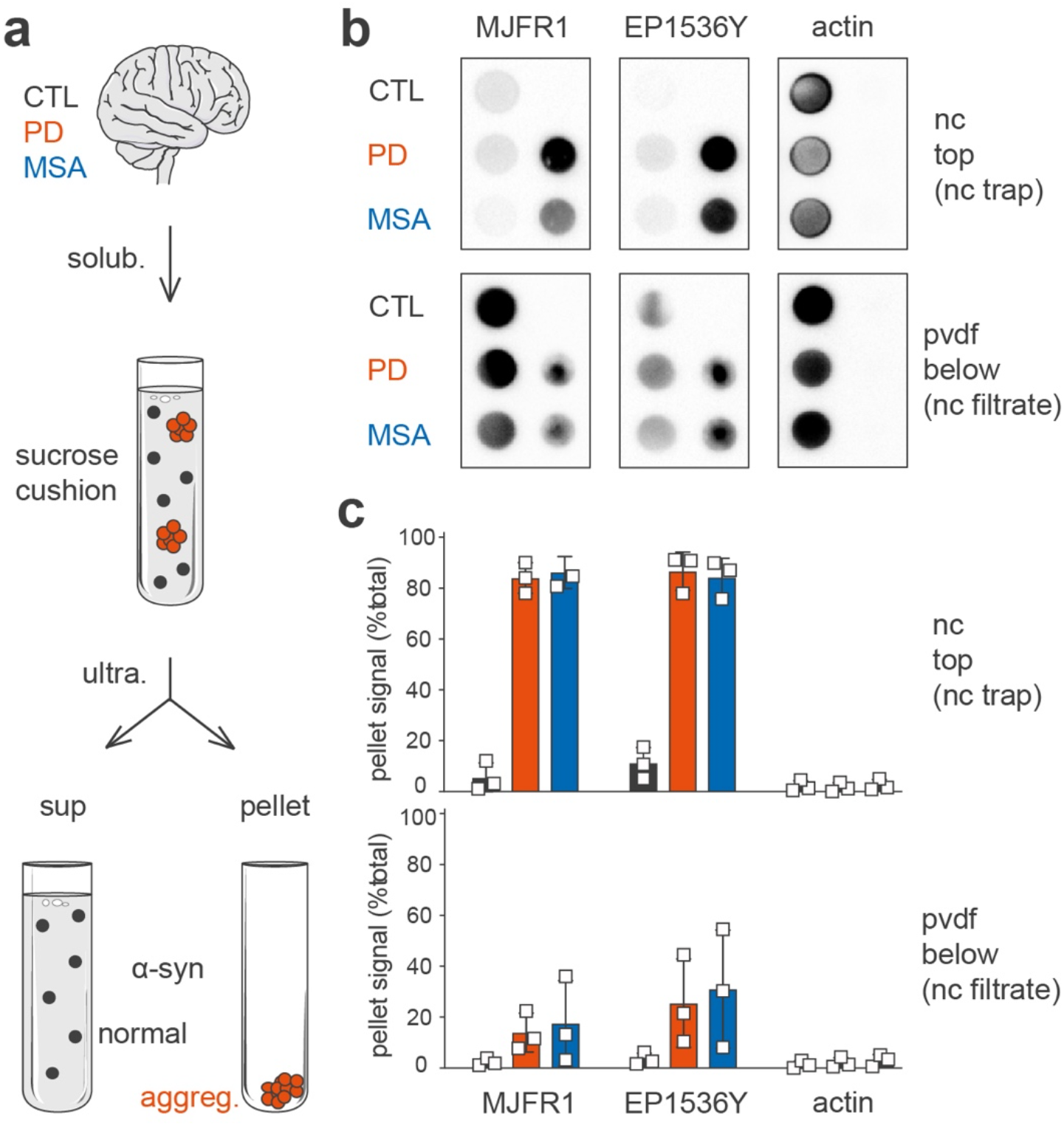
Extraction of pathological alpha-synuclein and separation from its normal counterparts from synucleinopathy brains by an adapted Sarkospin procedure. (**a**) Schematic representation of the adapted Sarkospin protocol. Human control, PD, or MSA brain samples were homogenized and solubilized with sarkosyl at 37°C in the presence of Benzonase before a single ultracentrifugation step within a sucrose cushion to separate supernatant and pellet fractions. (**b-c**) Biochemical analysis of Sarkospin fractions extracted from human brain samples. Supernatants and pellets were subjected to filter trap on layered nitrocellulose (nc, top) and pvdf (bottom) membranes immunoblotted against human α-synuclein (MJFR1), S129-phosphorylated α-synuclein (EP1536Y) or ß-actin (actin). (**b**) Representative pictures of the immunoblots for all samples, membranes, and antibodies. (**c**) Quantifications of each protein’s percentage residing in Sarkospin supernatant (light) or pellet (dark) for all human samples used in this experiment. Bars represent mean with standard deviations for n=3 biologically independent human brain samples per group.

The results indicate that the adapted Sarkospin procedure efficiently separated pathological aggregated α-syn from its soluble counterparts. As observed in Fig. 1b, high molecular weight α-syn species that are filter-trapped on nitrocellulose membrane are specifically present in the pellet fractions of PD and MSA samples (MJFR1, nc). In contrast, small mono- and oligomeric forms of the protein that get to pvdf membranes are observable mainly in all groups’ supernatants (MJFR1, pvdf).

Another marker of pathological forms of the protein, namely its S129-phosphorylation (EP1536Y positive signal), indicates that large insoluble pS129-α-syn-containing aggregates are isolated in the pellet fractions for synucleinopathy samples (EP1536Y, nc) as opposed to the physiological EP1536Y-positive soluble entities found in all supernatants (EP1536Y, pvdf). Also, α-syn aggregates purified in synucleinopathy sample pellets have proved to be peculiarly resistant to denaturation (SDS, heat) and proteolysis (Proteinase K) while supernatants comprised only sensitive monomeric bands (Sup. Fig. 2a, b). Notably, actin, yet physiologically forming large assemblies, is found only in supernatant Sarkospin fractions and witnesses the procedure’s efficient solubilization (Fig. 1a, b).

By quantification on filter-trap immunoblots for n=9 brains, of Sarkospin supernatant and pellet fractions content in the proteins mentioned above, it appeared that over 80% of the nc-trapped α-syn in brain extracts from PD and MSA patients was pelleted, and less than 5% in brain extracts from control subjects (Fig. 1c, MJFR1 nc). In PD and MSA samples, S129-phosphorylated α-syn coincided with the insoluble α-syn pool since 90% of the pS129-α-syn signal (Fig. 1c, EP1536Y nc) was found in the pellets. In clear contrast, this figure was less than 5% for the control subjects. Noteworthy, a small subset of insoluble aggregates passes to pvdf membranes, but the majority (60-70% for PD and MSA, 99% for CTL) of α-syn present on this support is from supernatants.

Collectively, these results indicate that the novel Sarkospin procedure efficiently extracts pathological α-syn aggregates from human brain samples and physically separates these entities from physiological mono- and oligomeric α-syn forms present in healthy brain samples. Thus, we attained a purification method compatible with a subsequent proteomic analysis to extract and identify insoluble proteins from synucleinopathy subject brains and their direct comparison with parallel Sarkospin extractions from control patient brains.

### Prime enrichment in alpha-synuclein and unidentified insoluble proteins in PD and MSA Sarkospin pellet fractions

Proteomic analysis of Sarkospin pellets extracted from control, PD, and MSA samples (n=3 independent subject brains per group) identified a total of 1022 proteins that resisted sarkosyl solubilization (Fig. 2a, b, Appendix A, Master Protein). The quantifications and comparisons of each protein’s normalized abundance in synucleinopathy vs. control pellets allowed the calculation of individual fold change (FC) and statistical significance (p-value of multiple t-tests) of the difference of abundance between groups. The results show the gating of 130 and 160 proteins found significantly enriched (FC > 1.5 and p<0.05) in PD (Fig. 2a, red) and MSA (Fig. 2b, blue) vs. control, respectively (Appendix B-C). Impressively, α-syn is by far the most significantly enriched protein in all synucleinopathy samples, with a fold change of disease vs. control close to 40 while the second most enriched protein reaches an FC of 10 (Fig. 2a, b, and Appendix B-C, α-syn: PD vs. CTL FC=36 p=7.0×10^-3^; MSA vs. CTL FC=38 p=3.5×10^-3^).

**Fig. 2.**
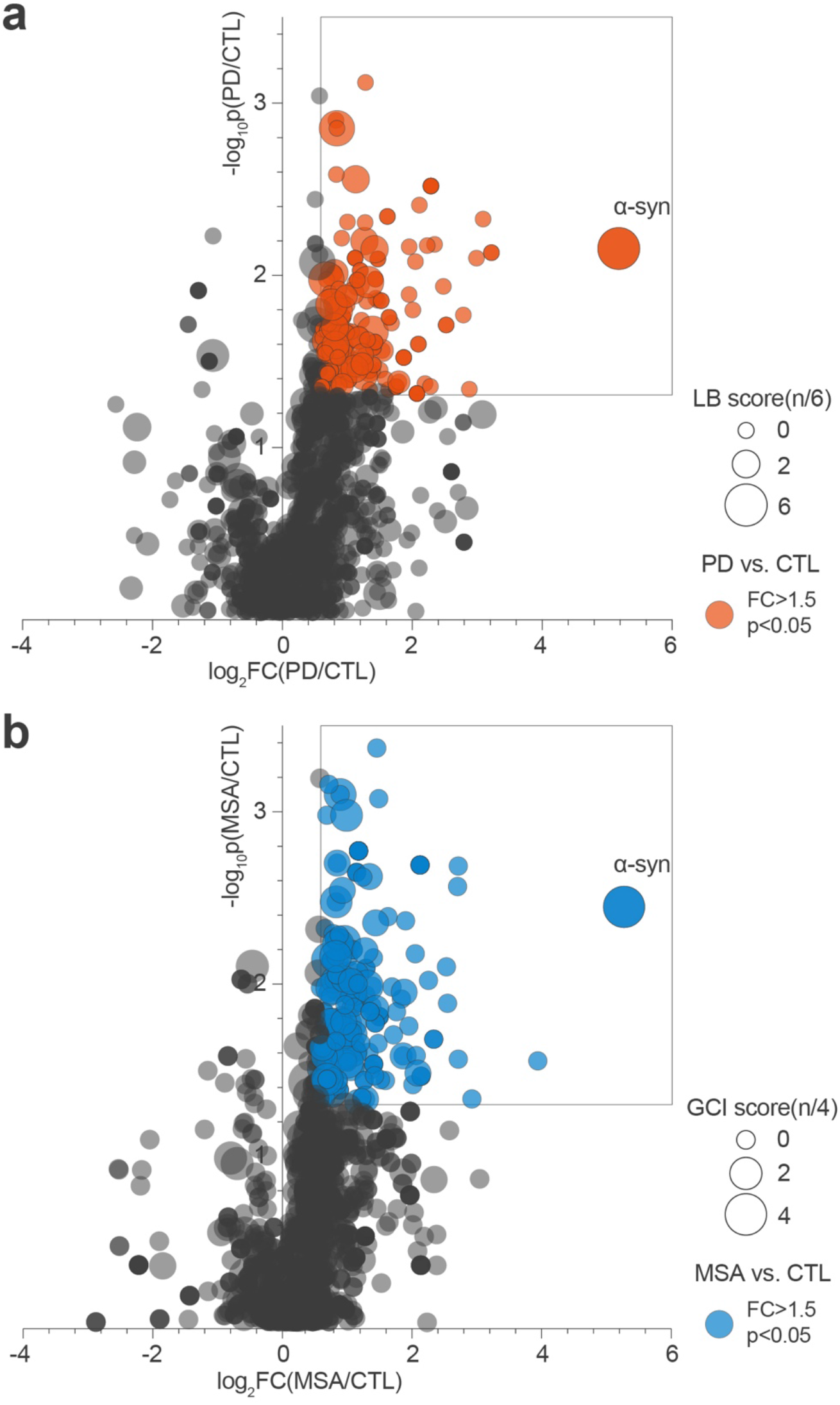
Prime enrichment in alpha-synuclein and unidentified insoluble proteins in PD and MSA Sarkospin pellet fractions. Bubble volcanos charts representing mass spectrometry analysis of the protein content of all n=9 pellets. Each of the 1022 master proteins identified in the present proteomic study is plotted as a circle positioned upon the fold change (log_2_FC) and statistical significance (-log_10_pValue) of its enrichment in PD (**a**) and MSA (**b**) insoluble pellets as compared to controls, respectively. Fold changes represent ratios of normalized abundance of a protein (normalized between individuals to the abundance of all proteins) between disease and control groups, with p values of the respective multiple t-tests ran on n=3 biologically independent human brain samples per group. Colored bubbles indicate proteins significantly enriched in disease (red for PD, blue for MSA) vs. control pellets, with a FC>1.5 and p<0.05. A total of 206 are found significantly enriched in any of the two synucleinopathy groups, with 130 proteins for PD (a, red) and 160 for MSA (b, blue). Bubbles are sized upon their respective LB or GCI scores from mass spectrometry analysis in the literature. These scores refer to the number of articles (6 for LBs, 4 for GCIs) identifying this specific protein in the respective pathological inclusions out of ten articles^3,5,6,20,21,23–25,27^.

We calculated bibliographic LB and GCI scores for the identification in this dataset of the proteins previously detected by proteomic analysis in the literature as components of pathological cytoplasmic inclusions. These indices relate to the number of mass spectrometry studies identifying the indicated protein in LBs and GCIs out of six^3,6,23–25,27^ and four^5,6,23,25^ publications, respectively.

Interestingly many of the most significantly enriched candidates were not identified in previous studies and had a null bibliographic LB and GCI score (smallest-sized red and blue bubbles close to α-syn on Fig. 2a and 2b, respectively). Besides, many proteins previously identified (i.e., with high bibliographic LB or GCI indices) are found in the extracts. Still, they are not significantly enriched in synucleinopathy pellets compared to control patient samples with no synucleinopathy (large dark bubbles on Fig. 2a, b). This could be explained by the absence in previous studies of a comparator group with equivalent fractions from healthy brains allowing to rule out an unselective enrichment. Indeed, for proteins physiologically highly expressed in the brain, their high abundance could simply lead to an increased detection probability whatever the IB purification method, irrespective of the IB pathology.

We challenged this possibility for all the 1022 proteins present in the insoluble proteome. Fig. 3 represents the absolute abundance of each protein in control and disease samples, with color-coding the score of identification in the literature (bibliographic LB and GCI scores, Fig. 3a, 3c) or in our proteomic analysis (Z score, Fig. 3b, 3d), and allows to highlight the association between these parameters. These results unambiguously show that most of the proteins in LB and GCI proteomes previously reported in the literature are abundant in both control and pathological insoluble fractions (colored “tip of the spear”, Fig. 3a, 3c). Instead, selecting the proteins showing a significant enrichment in IB vs. control brain fractions allows the identification of pathology-related proteins independently of their abundance level, as a result, even rare proteins can be detected as significantly enriched in PD or MSA pellets as compared to controls (colored “side of the spear”, Fig. 3b, 3d).

**Fig. 3.**
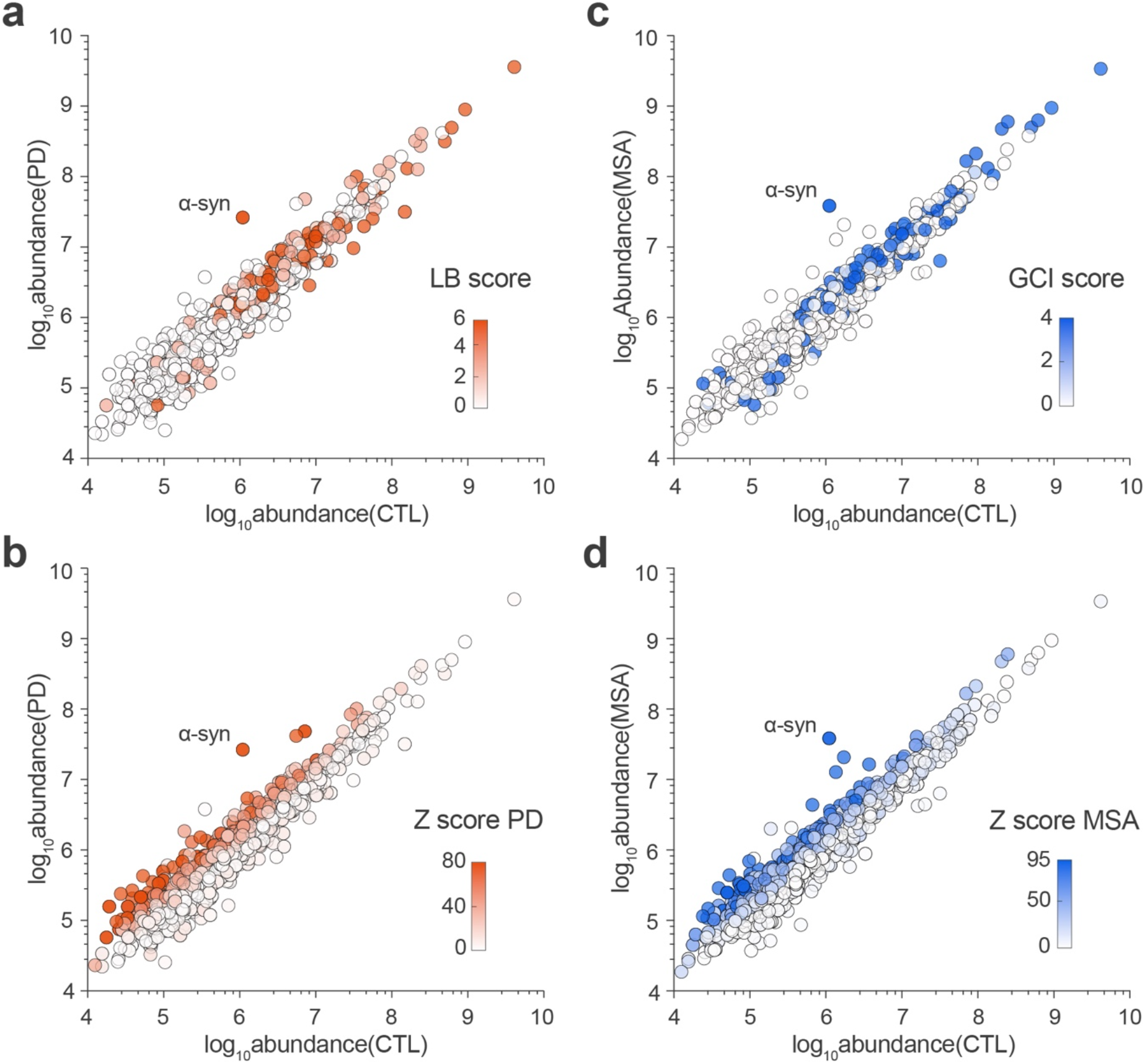
Identification and resolution of synucleinopathy insoluble proteomes independently of the physiological abundance of proteins. Spear charts representing mass spectrometry analysis of the protein content of all n=9 Sarkospin pellets. Each of the 1022 master proteins identified in the present proteomic study is plotted as a circle positioned upon the total abundance of the protein (log_10_Abundance, A.U.) in control samples and its total abundance in PD (**a-b**) and MSA (**c-d**) samples. (**a**) and (**c**) spear charts are respectively color-coded upon the protein LB and GCI scores described in Fig. 2. (**b**) and (**d**) spear charts are color-coded respectively upon the protein Z score for PD and MSA, with Z = FC x (-log_10_ p) for each disease group.

Therefore, the present Sarkospin procedure, coupled with a subsequent comparative proteomic analysis, is an adequate method for unraveling insoluble proteomes and unbiasedly refining the list of candidates by considering their fold enrichment with regards to control samples. This approach defines the insoluble proteomes specific for each synucleinopathy and identifies LB and GCI components.

### Major overlap between PD and MSA insoluble proteomes

A total of 206 unique proteins were gated previously as significantly enriched in at least one of the synucleinopathy types compared to controls (Fig. 2a and 2b, red and blue, Appendix D). Among this group of proteins, 84 are significantly enriched in PD as well as in MSA compared to controls with FC>1.5 and p<0.05 (Appendix E). Inversely, among the 206 disease-associated proteins, only seven proteins were found to be selectively enriched in one synucleinopathy compared to the other, with three PD-specific and four MSA-specific (Fig. 4, red and blue, Appendix F-G). This highlights the extreme narrowness of the PD- and MSA-specific insoluble proteomes and indicates that the most substantial part of synucleinopathy-associated insoluble proteomes is shared between these two pathologies.

**Fig. 4.**
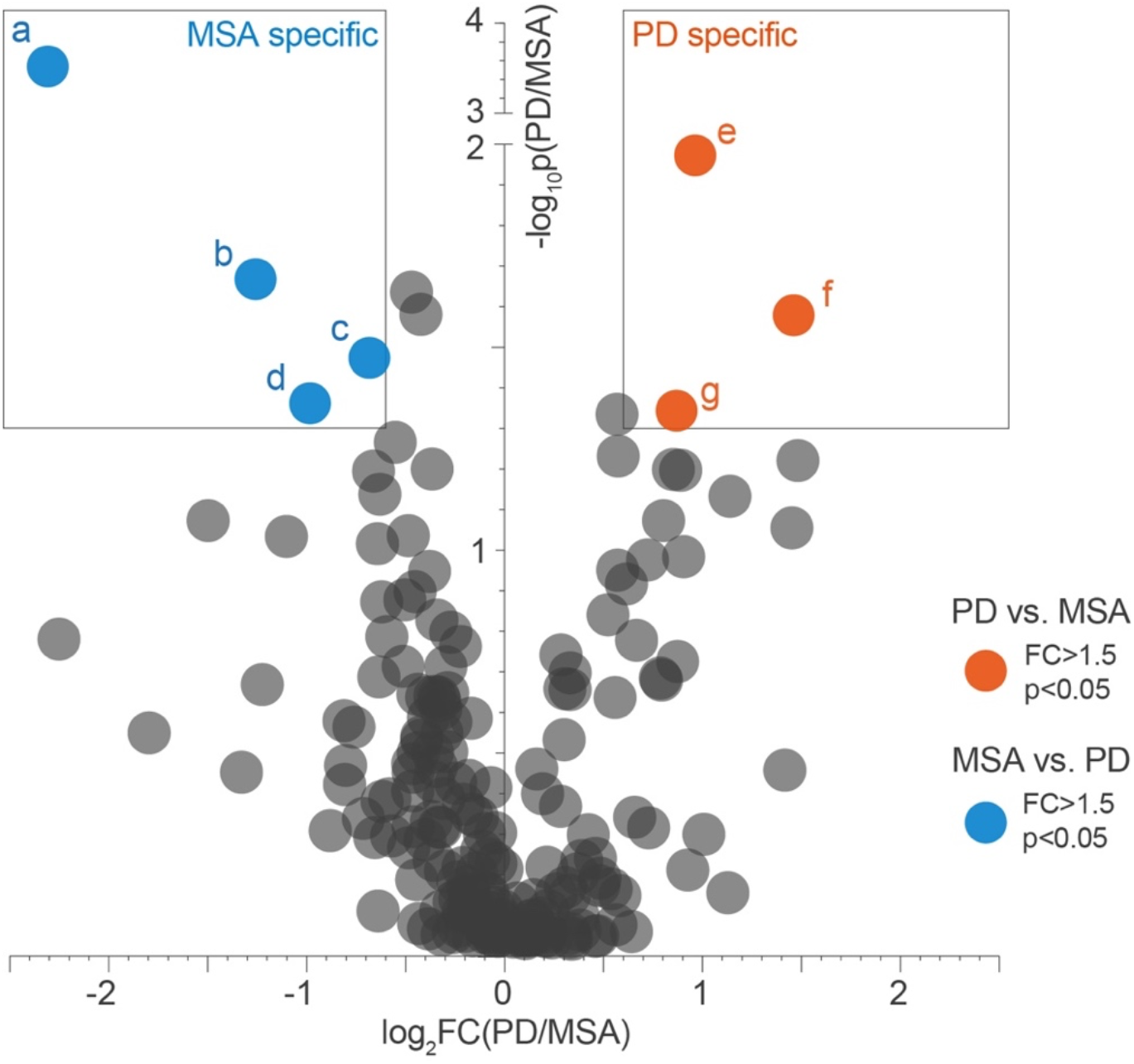
The remarkable overlap between PD and MSA respective insoluble proteomes. The bubble chart represents mass spectrometry analysis of the 206 proteins gated in Fig. 2 that are significantly enriched in PD or MSA compared to control Sarkospin pellets. Each protein is positioned upon the fold change (log_2_FC) of its enrichment in PD insoluble pellets compared to MSA, and the statistical significance of the protein abundance difference between PD and MSA Sarkospin pellets (-log_10_ p). Color-coding corresponds to each protein’s gating: red: n=3 proteins significantly enriched in PD and blue: n=4 proteins significantly enriched in MSA pellets as opposed to the other disease group respectively, and black: n=199 proteins not significantly different between PD and MSA. Significantly enriched terms a FC > 1.5 with p < 0.05 with n=3 independent biological samples per group. **a:** Protein phosphatase methylesterase 1; **b:** Vesicle-fusing ATPase; **c:** Calnexin; **d:** Phospholipid – transporting ATPase IA; **e:** Collagen alpha-1(XVIII) chain; **f:** 39S ribosomal protein L47, mitochondrial; **g:** Receptor expression-enhancing protein 1.

In other words, these data show the tremendous overlap of PD and MSA-associated insoluble proteomes. More specifically, in the most enriched proteins for both diseases or even for MSA specifically, a consistent proportion of candidates have been identified to play a role in PD pathogenesis (Appendix C, E, G). Namely, for example, NipSnap-1 (average FC=7 p=8.0×10^-3^) was shown to be implicated in Parkin-related mitophagy^30^; Septin-5 (average FC=4.6 p=2.5×10^-3^) was associated with dopamine-dependent neurotoxicity in early-onset PD also linked to Parkin^31^; even Protein phosphatase methylesterase 1 which we find to be MSA-specific (Fig. 4, a) has previously been also implicated in PD pathogenesis as enriched in PD subject substantia nigra^32^ and playing a role in α-syn phosphorylation through PP2A activity regulation^33^.

Notably, from the four MSA-specific candidates we identify, none shows a glial-specific expression or a glial tropism (Fig. 4, a-d, Appendix G). Besides protein phosphatase methylesterase 1 described above (Fig. 4, a), Phospholipid transporting ATPase IA acts as amino-phospholipid translocase at the neuronal plasma membrane (Fig. 4, d); Calnexin plays a role in receptor-mediated endocytosis at the synapse (Fig. 4, c); and Vesicle-fusing ATPase is directly implicated in vesicle-mediated transport and fusion of vesicles to membranes (Fig. 4, b).

Altogether, while the absence of oligodendroglial makers in the MSA-specific insoluble proteome is unexpected, the massive overlap of PD and MSA insoluble proteomes could suggest a common neuronal origin of these insoluble entities in both PD and MSA.

To put this possibility under scrutiny, we compared the lists of 130 and 160 proteins enriched respectively in PD and MSA pellets, to the published insoluble proteome found in an experimental neuronal synucleinopathy modelized in mouse neuronal cultures using preformed α-syn fibrils (PFFs) as seeds^12^ (Sup. Fig. 3). Strikingly, 38 and 58 insoluble proteins are shared between these PFF-treated neurons and PD and MSA respectively, with several – such as NipSnap-1 for PD, and Arginase-1 for MSA, showing a substantial enrichment in both proteomics studies (Sup. Fig. 3a and 3B). This striking overlap with yet completely independent studies and samples types validates several candidates, and brings a strong support to the notion that GCIs in MSA are made of α-syn amyloid building blocks preassembled in neurons

To confirm the enrichment of these candidates in PD, MSA or PFF-treated neurons insoluble proteomes, we ran an independent Sarkospin assay and subjected the fractions to the dot blot analysis used for the validation of Sarkospin (Fig. 1), with antibodies directed to ten proteins of interest, after validating their specificity on western blots (Sup. Fig. 4 and 5, Methods section). The latter analysis allowed to eventually validate the significant enrichment of proteins such as NipSnap-1 and Septin-5 in PD pellets or Arginase-1, Fumarase, and ATP8A1 in MSA pellets.

### Gene ontology analysis is supportive of the prime neuronal origin of GCIs and indicative of the absence of cross-seeded amyloid partners like tau or TDP-43

To better decipher the cellular localization or the implication in specific pathways and processes of the proteins identified in pathological insoluble proteomes, possibly indicating their origin and reason of enrichment, we ran a gene ontology (GO) analysis. The results of the GO overrepresentation test were performed on the list of 199 overlap proteins (Fig. 4, black) compared to the list of all 1022 proteins identified. This statistical test allows identifying clusters of biological processes, molecular functions, and cellular components specifically overrepresented in synucleinopathy insoluble proteomes by quantifying their respective enrichment and significance compared to controls (Fig. 5, fold change, p-value, Z score and #proteins, Appendix H).

**Fig. 5.**
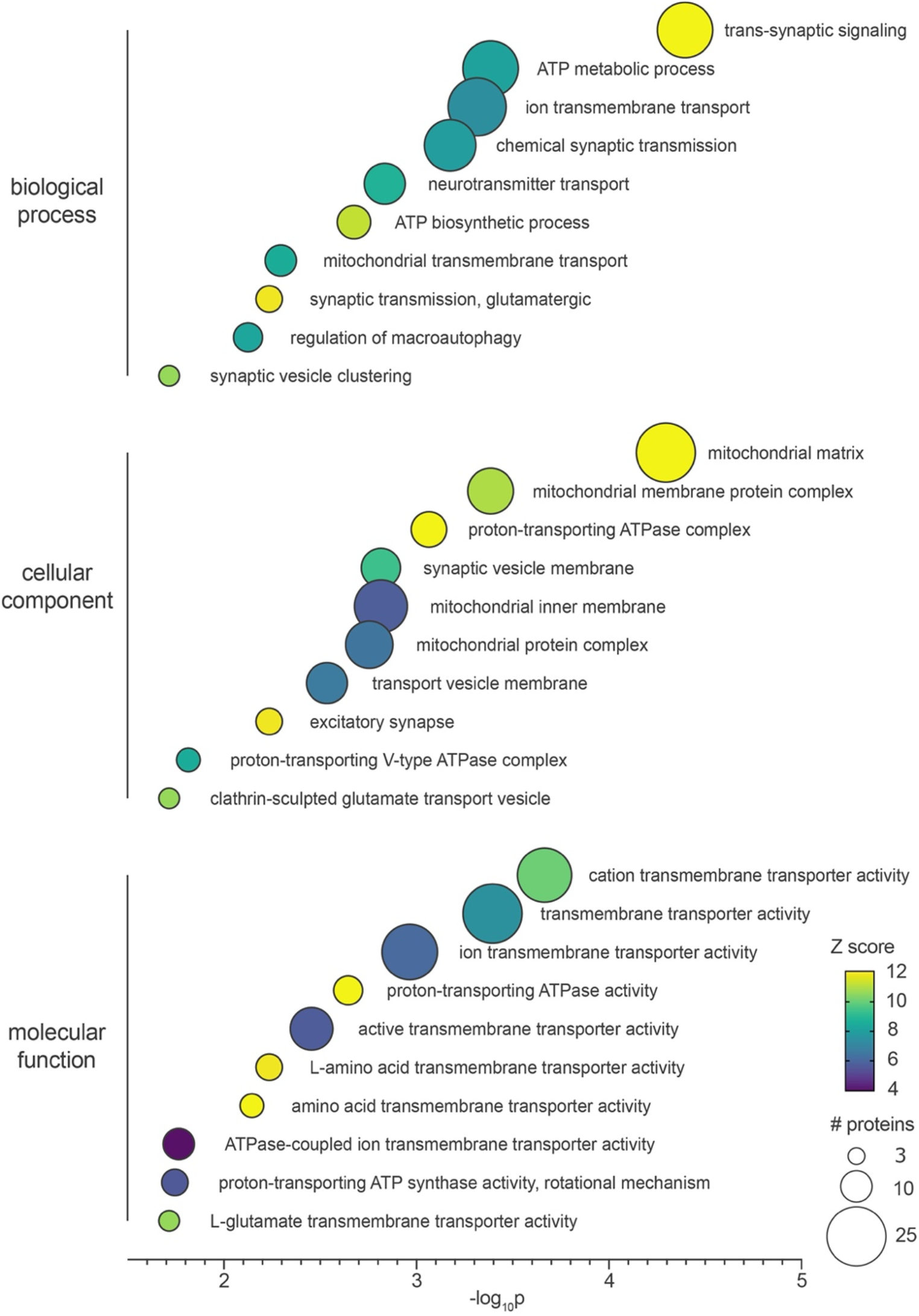
Gene ontology analysis of synucleinopathy-associated insoluble proteomes indicates a plausible neuronal origin of GCIs. The list of 199 proteins gated previously (206 proteins enriched in both synucleinopathies −7 specific to PD or MSA, Fig. 4 black, Appendix D-F-G) was compared to the total list of 1022 proteins found in all Sarkospin pellets (Appendix A) using Panther GeneOntology to calculate the statistical overrepresentation of biological processes (top), cellular components (middle) and molecular functions (bottom) gene ontology clusters in the synucleinopathy protein list. The bubble chart is representing the statistical significance (-log_10_p, multiple comparisons corrected binomial test) of gene ontology clusters, with their color-coded Z score (FC x (-log_10_p)) and size-coded number of represented proteins. Only significantly enriched (FC > 1.5; p < 0.05; # proteins > 3) clusters are represented on the graph (Appendix H).

First of all, a clear-cut result is that no cross-seeded amyloidogenic partner proteins are associated with fibrillar α-syn in PD and MSA.

The bubble chart representation of GO analysis in Fig. 5 compellingly shows that the most significantly enriched and importantly represented GO clusters for synucleinopathies indicate a neuronal imprint, with a remarkable synaptic and mitochondrial membrane trend (Fig. 5). Indeed, all biological processes, molecular functions, and cellular components reaching maximal fold change and statistical significance are related to synaptic and mitochondrial localization or pathways. Surprisingly, while even MSA-enriched insoluble proteomes are represented in these GO clusters, no protein identification nor GO cluster reflects a glial localization or tropism. Instead, all components of the latter clusters are enriched in neuronal cells. More specifically, mitochondrial actors (Mitochondrial proton/calcium exchanger protein, Fumarate hydratase, Pyruvate carboxylase, Sidoreflexin, Very long-chain specific acyl-CoA dehydrogenase, Cytochrome c oxidase) and even to a greater extend synaptic proteins (NipSnap-1, Septin-5, Synapsin-1, Synapsin-2, Synaptoporin, Glutamate receptor 2) are among the most enriched proteins in both PD and MSA pellets (Fig. 4).

Out of the 199 proteins used in the present GO analysis, 52 and 50 have been identified previously as components of LBs and GCIs, respectively, in at least one of the ten available proteomic studies^3,5,6,20,21,23–25,27^. Therefore, the insoluble proteomes of PD and MSA identified by Sarkospin most likely represent the core components of IBs. Their association with aggregated α-syn is tight enough to resist the harsh sarkosyl solubilization of the extraction procedure.

Collectively, the primary neuronal composition of synucleinopathy insoluble proteomes – and of MSA-specific one to a greater extend – the identification of PD-related proteins in MSA-enriched candidates and the synaptic origin of these components indicate the prime neuronal source of preassembled building blocks serving to the formation of oligodendroglial GCIs in MSA. Our conclusion is ideally in line with the experimental observation that in mice inoculated with α-syn PFFs, there is an initial florid neuronal synucleinopathy that develops to progressively decrease with time and leave the place to the progressive appearance of oligodendrocytes harboring GCI-like inclusions^34^. This is also in line with the notion that overexpression of α-syn in oligodendrocytes fails to cause the buildup of GCIs^28^. Finally, our observations do not support the idea that cross-seeding processes involving amyloidogenic proteins other than α-syn occur in PD and MSA.

## DISCUSSION

The development of a novel procedure by adaptation of our formerly published Sarkospin method^28,29^ (Fig. 1) allowed the extraction and purification of pathological aggregates from synucleinopathy subject brain tissues. The data presented here confirm that α-syn is the only amyloid and by far the most enriched protein present in PD and MSA sarkosyl-insoluble fractions of brain samples^6^ (Fig. 2). The presence of this neuronal protein in oligodendrocyte inclusions is the core point of the enigma we sought to address in the present study^35^: what are the origin and the reason for the formation of LBs and GCIs?

Characterizing the proteomes of LBs and GCIs has been repeatedly attempted in previous studies in order to understand which proteins are involved in their formation and their packing. Besides α-syn, mass spectrometry analysis of these IB-enriched fractions identified hundreds of proteins that potentially play a key role in the pathogenesis or the origin and mechanisms of formation of LBs and GCIs. These proteins include structural and cytoskeletal elements (neurofilaments, tubulins, tubulin polymerization promoting protein, TPPP/p25), α-synuclein-binding proteins (14-3-3, synphillin-1), components of the ubiquitin-proteasome system, α-ß-crystallin, heat shock proteins, or DJ-1. However, these candidates’ relevance to the molecular pathology of IBs can be limited by the ability of the experimental procedures to isolate IBs with sufficient purity and yield from the surrounding structures and protein contaminants. Recently, a purification method including a partial proteolysis step allowed the refinement of this list of proteins and demonstrated a high burden of synaptic vesicle-related proteins of the IBs^25^. These candidates are implicated in clathrin-mediated endocytosis (clathrin, AP-2 complex, dynamin), retrograde transport (dynein, dynactin, spectrin), or synaptic vesicle fusion (synaptosomal-associated protein 25, vesicle-associated membrane protein 2, syntaxin-1).

Using extracts from control, PD, and MSA brains, the comparative proteomic analysis we performed identified proteins enriched in PD and MSA brain samples (Fig. 2), with novel undescribed candidates, and refinement by excluding a list of previously identified proteins as probable false positives and/or protein not tightly associated with the sarkosyl-insoluble α-syn amyloids of IBs (Fig. 2)^3,5,6,20,21,23–25,27^.

The protein enrichment in the respective PD or MSA insoluble proteome showed a tremendous overlap between them, with only few proteins being significantly different (Fig. 4). Lastly, the latter results associated with a gene ontology analysis revealed compellingly that both insoluble proteomes are composed of the vast majority of neuronal proteins, with a significant synaptic overrepresentation (Fig. 5), as well as numerous mitochondrial proteins. These data are in total agreement with previous proteomic studies^25^ and microscopic analysis^4^ which revealed a burden of synaptic-vesicle-related proteins, and membranous structures, such as vesicles or dysmorphic mitochondria. Noteworthy, our study independently validated many synaptic proteins found in the previously mentioned study^25^. These results strengthen the hypothesis of the neuronal origin of GCIs. It is tempting to speculate that in MSA α-syn could primarily get aggregated in neurons, then released as preformed inclusions bodies, eaten-up by surrounding oligodendrocytes, and eventually matured/stored as inert GCIs^34^. The fact that oligodendrocyte-specific proteins are not enriched in MSA samples by our Sarkospin procedure is in line with the idea that the oligodendrocyte contribution is late, with glial proteins only secondarily and loosely associated with the GCIs and not properly with the amyloid core of the inclusion, previously pre-assembled in neurons.

Of particular importance, the list of proteins identified in the present study is restricted to the production of peptides identifiable by mass spectrometry, deriving from the presence of tryptic cleavage sites. Suppose one considers the high representation of neurodegenerative disease-associated proteins in partially trypsin-resistant proteins. In that case, it is possible that their enrichment here was underestimated, or even that some escaped identification. However, the conditions used here are known to reveal TDP-43 and tau^29^, and the absence of detection of cross-seeded amyloid partners does not thus derive from an insufficient proteolytic cleavage.

Extending the present study to other brain regions and at different pathology stages would be of particular interest. In pafticular, using amygdala or putamen extracts for focusing on LBs or GCIs protein compositions, respectively, should help refining the present lists and identifying less represented candidates. Also, Sarkospin extraction and purification of synucleinopathy subject brain tissue yet devoid of any inclusion^36^, and the insoluble proteomes’ comparison to the ones we defined here is of high importance for understanding the formation and composition of IBs. The Sarkospin procedure we developed here allows testing the seeding ability^37^ or the pathogenicity of aggregated proteins^38^, an asset to understand the role of the inclusions in the disease process. Also, facing these insoluble proteomes’ components to lists of identified interactants of mono-oligomeric physiological α-syn, or α-syn amyloid binders would help understand why the presence of these proteins in the inclusions and to decrypt their mechanisms of pathological association.

## METHODS

### Human brain samples

The samples were obtained from brains collected in a Brain Donation Program of the Brain Bank “GIE NeuroCEB” (Neuro-CEB BB-0033-00011). The consents were signed by the patients themselves or their next of kin in their name, in accordance with the French Bioethical Laws. The Brain Bank GIE NeuroCEB has been declared at the Ministry of Higher Education and Research and has received approval to distribute samples (agreement AC-2013-1887). Human cortices (cingulate gyrus) were dissected from freshly frozen post-mortem brain samples from n=3 control, sporadic PD or MSA subjects respectively. The choice of cingulate gyrus samples was based on a prior biochemical analysis of different brain regions from the CTL, PD and MSA subjects in this series by immunoblotting against pS129 phosphorylated (Sup. Fig. 1a) and aggregated (Sup. Fig. 1b) α-syn. On the basis of these data, we found that the cingulate gyrus was the most comparable region in terms of total amyloid α-syn load for these PD and MSA subjects. In addition, the pathological records regarding the contralateral hemispheres indicated the presence of substantial amounts of LBs and GCIs in this area for the PD and MSA subjects, respectively. We thus proceeded with a comparative proteomic analysis of the insoluble proteomes using this brain region.

### Human subject anonymized information

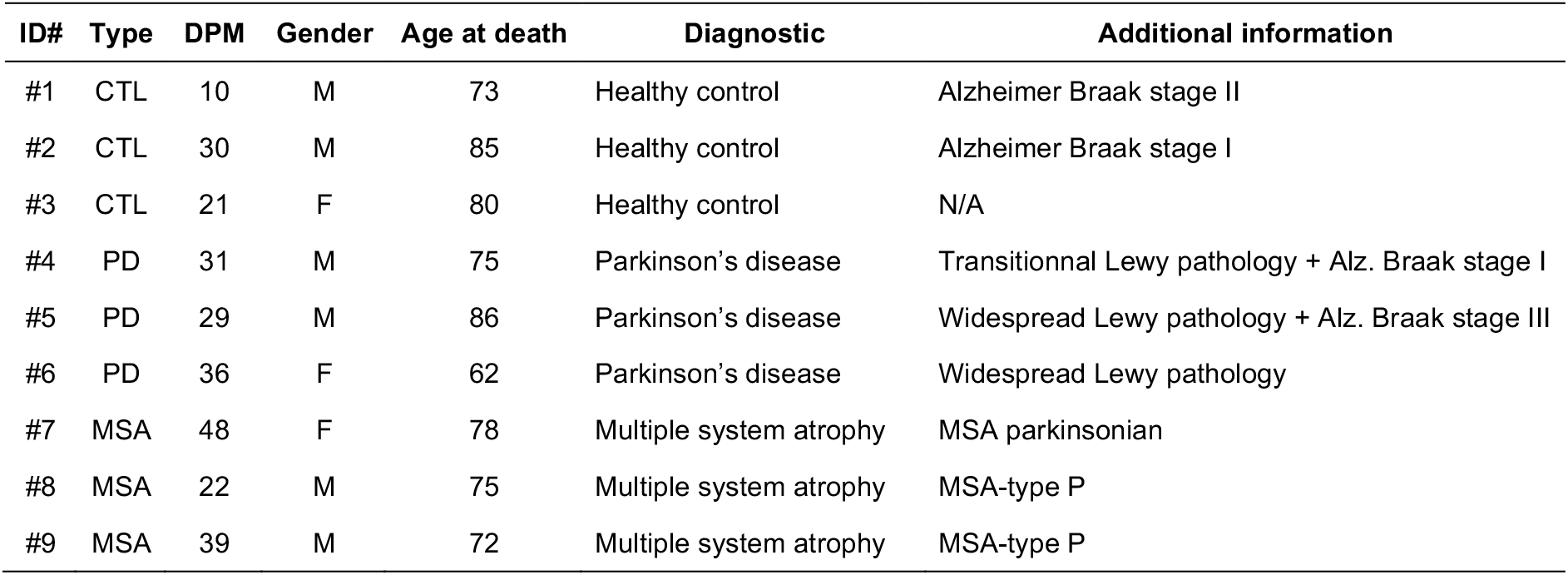

### Sarkospin procedure

For the extraction and purification of aggregates from brains samples, the Sarkospin procedure was adapted from previously published protocols^28,29^. Brain tissue samples were homogenized at 10% (w/v) in solubilization buffer (SB): 10 mM Tris pH 7.5, 150 mM NaCl, 0.1 mM EDTA, 1 mM DTT, Complete EDTA-free protease inhibitors (Roche) and PhosSTOP phosphatase inhibitors (Roche) using a gentleMACS Octo Dissociator (Miltenyi Biotec) with M Tubes, and the Protein extraction program. Samples were mixed 1:1 with SB 4% (w/v) N-lauroyl-sarcosine (sarkosyl, Sigma), 2 U.μl^-1^ Benzonase (Novagen) and 4 mM MgCl2, reaching a final volume of 500 μl. Sarkospin solubilization was then performed by incubating the samples at 37 °C under constant shaking at 600 rpm (Thermomixer, Eppendorf) for 45 min. Solubilized samples were then mixed 1:1 with SB 40% (w/v) sucrose, without sarkosyl, MgCl_2_ or Benzonase, in 1 ml polycarbonate ultracentrifuge tubes (Beckman Coulter) and centrifuged at 250,000 g for 1 hour at room temperature with a TLA 120.2 rotor using an Optima XP benchtop ultracentrifuge (Beckman Coulter). Supernatant were collected by pipetting. For filter trap and SDS-PAGE immunoblot analysis, pellets were resuspended directly in the tube with 100 μL of the buffer corresponding to the supernatant (SB 1% sarkosyl 20% sucrose), and mixed with the same buffer in a fresh tube for reaching 1ml (equal volumes to supernatant). For proteomics analysis, pellets were resuspended in 100 μL SB, and equalized for their total protein concentration quantified by bicinchoninic acid (BCA) assay (Thermo), prior to denaturation in Laemmli buffer.

### Analysis of the protein contents of Sarkospin fractions by filter trap

For filter trap assays, native Sarkospin fractions were spotted onto layered nitrocellulose and PVDF 0.2 μm membranes (Protran, GE) using a dot blot vacuum device (Whatman). Membranes were fixed for 30 min in PBS with PFA 0.4% (v/v) (Sigma) final concentration. After three washes with PBS, membranes were blocked with 5% (w/v) skimmed powder milk in PBS-Tween 0.5% (v/v) and probed with primary and secondary antibodies in PBS-Tween with 4% (w/v) BSA (Antibody Table). Immunoreactivity was whether visualized by chemiluminescence or infrared using Clarity ECL and Chemidoc (Biorad) or Odissey systems (Li-Cor) respectively.

### Proteinase K treatments and western blot

For PK resistance assays, equal volumes of solubilized homogenates, Sarkospin supernatants or pellets fractions were treated or not with 1 μg.ml^-1^ Proteinase K (Sigma) from 0 to 60 minutes at 37°C. At the end of the indicated time, samples were added Laemmli 1x prior to denaturation at 95 °C for 5 min, and loaded on Mini-Protean TGX 12% gels (Biorad) followed by SDS-PAGE electrophoresis. Gels were whether stained for total protein amount (silver stain, Biorad) or transferred on nitrocellulose 0.2 μm membranes with Trans-Blot Turbo transfer system (Biorad) using the Mixed molecular weight program. Membranes were fixed with PFA, and proteins were immunolabelled as described for filter trap.

### Mass spectrometry analysis of Sarkospin pellets

The mass spectrometry proteomics data have been deposited to the ProteomeXchange Consortium via the PRIDE partner repository with the dataset identifier **PXD024998**.

### Sample preparation and protein digestion

Protein samples were solubilized in Laëmmli buffer and 10 μg per sample were deposited onto SDS-PAGE gel for concentration and cleaning purpose. Separation was stopped once proteins have entered resolving gel. After colloidal blue staining, bands were cut out from the SDS-PAGE gel and subsequently cut in 1 mm x 1 mm gel pieces. Gel pieces were destained in 25 mM ammonium bicarbonate 50% acetonitrile (ACN), rinsed twice in ultrapure water and shrunk in ACN for 10 min. After ACN removal, gel pieces were dried at room temperature, covered with the trypsin solution (10 ng.μl^-1^ in 50 mM NH_4_HCO_3_), rehydrated at 4 °C for 10 min, and finally incubated overnight at 37 °C. Spots were then incubated for 15 min in 50 mM NH_4_HCO_3_ at room temperature with rotary shaking. The supernatant was collected, and an H_2_O/ACN/HCOOH (47.5:47.5:5) extraction solution was added onto gel slices for 15 min. The extraction step was repeated twice. Supernatants were pooled and dried in a vacuum centrifuge. Digests were finally solubilized in 0.1% HCOOH.

### nLC-MS/MS and label-free quantitative data analysis

Peptide mixture was analyzed on an Ultimate 3000 nanoLC system (Dionex, Amsterdam, The Netherlands) coupled to an Electrospray Orbitrap Fusion™ Lumos™ Tribrid™ Mass Spectrometer (Thermo Fisher Scientific, San Jose, CA). Ten microliters of peptide digests were loaded onto a 300 μm inner diameter x 5 mm C_18_ PepMap™ trap column (LC Packings) at a flow rate of 10 μL.min^-1^. The peptides were eluted from the trap column onto an analytical 75 mm id x 50 cm C18 Pep-Map column (LC Packings) with a 4-40% linear gradient of solvent B in 105 min (solvent A was 0.1% formic acid and solvent B was 0.1% formic acid in 80% ACN). The separation flow rate was set at 300 nL.min^-1^. The mass spectrometer operated in positive ion mode at a 1.8 kV needle voltage. Data were acquired using Xcalibur 4.1 software in a data-dependent mode. MS scans *(m/z* 375-1500) were recorded at a resolution of R = 120 000 (@ m/z 200) and an automated gain control target of 4 x 10^5^ ions collected within 50 ms. Dynamic exclusion was set to 60 s and top speed fragmentation in Higher-energy collisional dissociation (HCD) mode was performed over a 3 s cycle. MS/MS scans with a target value of 3 x 10^3^ ions were collected in the ion trap with a maximum fill time of 300 ms. Additionally, only +2 to +7 charged ions were selected for fragmentation. Others settings were as follows: no sheath nor auxiliary gas flow, heated capillary temperature, 275 °C; normalized HCD collision energy of 30% and an isolation width of 1.6 m/z. Monoisotopic precursor selection (MIPS) was set to Peptide and an intensity threshold was set to 5 x 10^3^.

### Database search and results processing

Data were searched by SEQUEST through Proteome Discoverer 2.3 (Thermo Fisher Scientific Inc.) against the *Homo sapiens* Reference Proteome Set (from Uniprot 2019-03; 73,645 entries). Spectra from peptides higher than 5000 Da or lower than 350 Da were rejected. The search parameters were as follows: mass accuracy of the monoisotopic peptide precursor and peptide fragments was set to 10 ppm and 0.6 Da respectively. Only b-and y-ions were considered for mass calculation. Oxidation of methionines (+16 Da) and protein N-terminal Acetylation (+42 Da) were considered as variable modifications and carbamidomethylation of cysteines (+57 Da) as fixed modification. Two missed trypsin cleavages were allowed. Peptide validation was performed using Percolator algorithm^39^ and only “high confidence” peptides were retained corresponding to a 1% False Positive Rate at peptide level. Peaks were detected and integrated using the Minora algorithm embedded in Proteome Discoverer. Proteins were quantified based on unique peptides intensities. Normalization was performed based on total protein amount. Protein ratio were based on protein abundances calculated as the sum of the three most intense peptides. Two-ways ANOVA tests were calculated considering the three values of protein abundance in each comparison. Quantitative data were considered for proteins quantified by a minimum of 2 peptides, fold changes above 1.5 and a statistical p-value lower than 0.05.

### Gene Ontology analysis

Gene ontology analysis was performed online using Panther Geneontology website: http://pantherdb.org. Lists of Uniprot accession IDs of the different gated proteins (PD and MSA groups) were used as sample Homo sapiens list, and were searched for statistical overrepresentation test against the list of all proteins identified in the present MS study, with biological process, molecular function and cellular components annotations. Results of multiple comparison corrected t-tests gave the indicated fold enrichment, number of proteins, and p value for the overrepresentation of each gene ontology cluster identified. Z scores were calculated as Z = fold change x (-log_10_(p value)).

### Antibodies used in this study

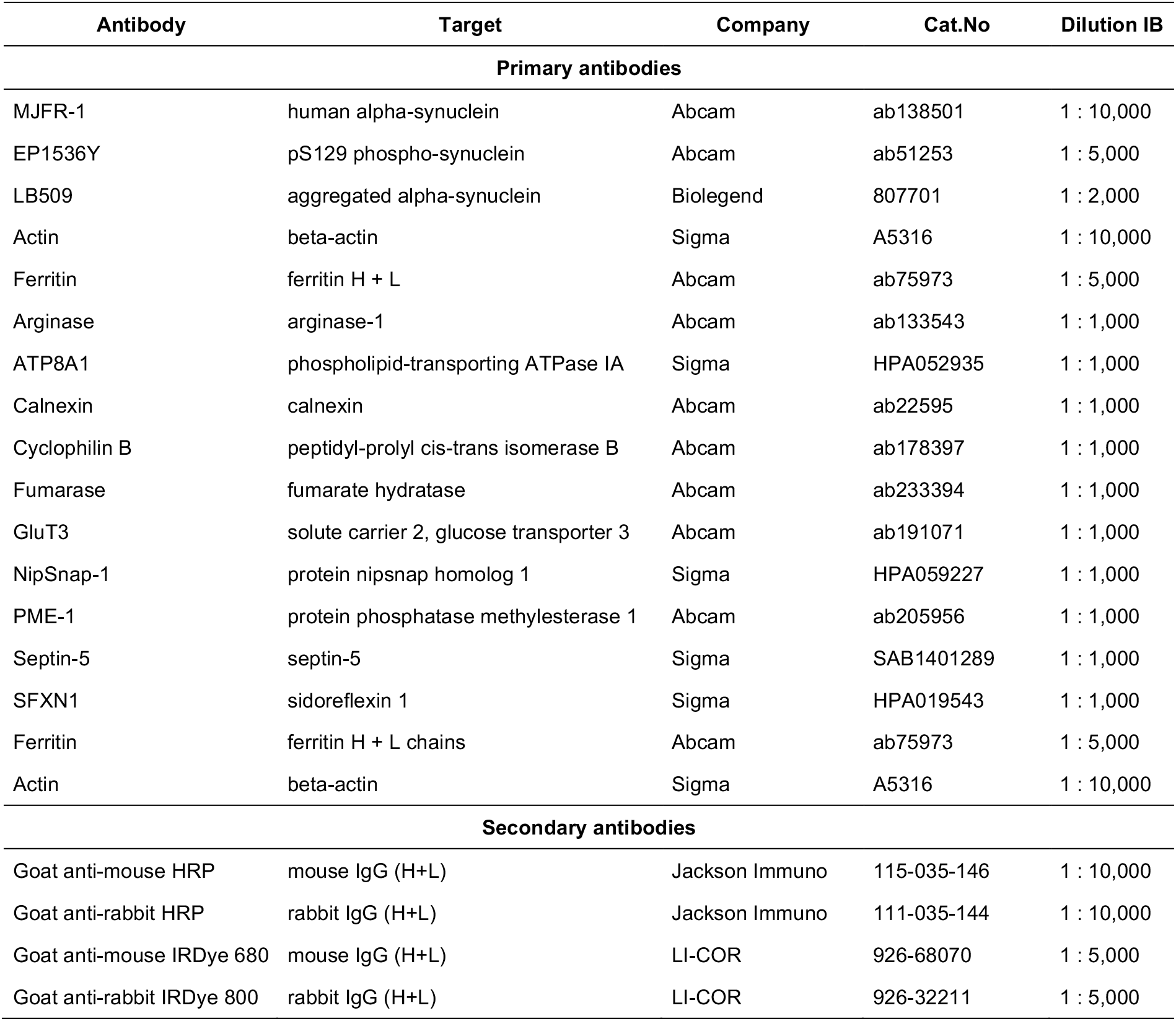

| Antibody | Target | Company | Cat.No | Dilution IB |
| --- | --- | --- | --- | --- |
| **Primary antibodies** |  |  |  |  |
| MJFR-1 | human alpha-synuclein | Abcam | ab138501 | 1 : 10,000 |
| EP1536Y | pS129 phospho-synuclein | Abcam | ab51253 | 1 : 5,000 |
| LB509 | aggregated alpha-synuclein | Biolegend | 807701 | 1 : 2,000 |
| Actin | beta-actin | Sigma | A5316 | 1 : 10,000 |
| Ferritin | ferritin H + L | Abcam | ab75973 | 1 : 5,000 |
| Arginase | arginase-1 | Abcam | ab133543 | 1 : 1,000 |
| ATP8A1 | phospholipid-transporting ATPase IA | Sigma | HPA052935 | 1 : 1,000 |
| Calnexin | calnexin | Abcam | ab22595 | 1 : 1,000 |
| Cyclophilin B | peptidyl-prolyl cis-trans isomerase B | Abcam | ab178397 | 1 : 1,000 |
| Fumarase | fumarate hydratase | Abcam | ab233394 | 1 : 1,000 |
| GluT3 | solute carrier 2, glucose transporter 3 | Abcam | ab191071 | 1 : 1,000 |
| NipSnap-1 | protein nipsnap homolog 1 | Sigma | HPA059227 | 1 : 1,000 |
| PME-1 | protein phosphatase methylesterase 1 | Abcam | ab205956 | 1 : 1,000 |
| Septin-5 | septin-5 | Sigma | SAB1401289 | 1 : 1,000 |
| SFXN1 | sidoreflexin 1 | Sigma | HPA019543 | 1 : 1,000 |
| Ferritin | ferritin H + L chains | Abcam | ab75973 | 1 : 5,000 |
| Actin | beta-actin | Sigma | A5316 | 1 : 10,000 |
| **Secondary antibodies** |  |  |  |  |
| Goat anti-mouse HRP | mouse IgG (H+L) | Jackson Immuno | 115-035-146 | 1 : 10,000 |
| Goat anti-rabbit HRP | rabbit IgG (H+L) | Jackson Immuno | 111-035-144 | 1 : 10,000 |
| Goat anti-mouse IRDye 680 | mouse IgG (H+L) | LI-COR | 926-68070 | 1 : 5,000 |
| Goat anti-rabbit IRDye 800 | rabbit IgG (H+L) | LI-COR | 926-32211 | 1 : 5,000 |

### Chemicals and kits used in this study

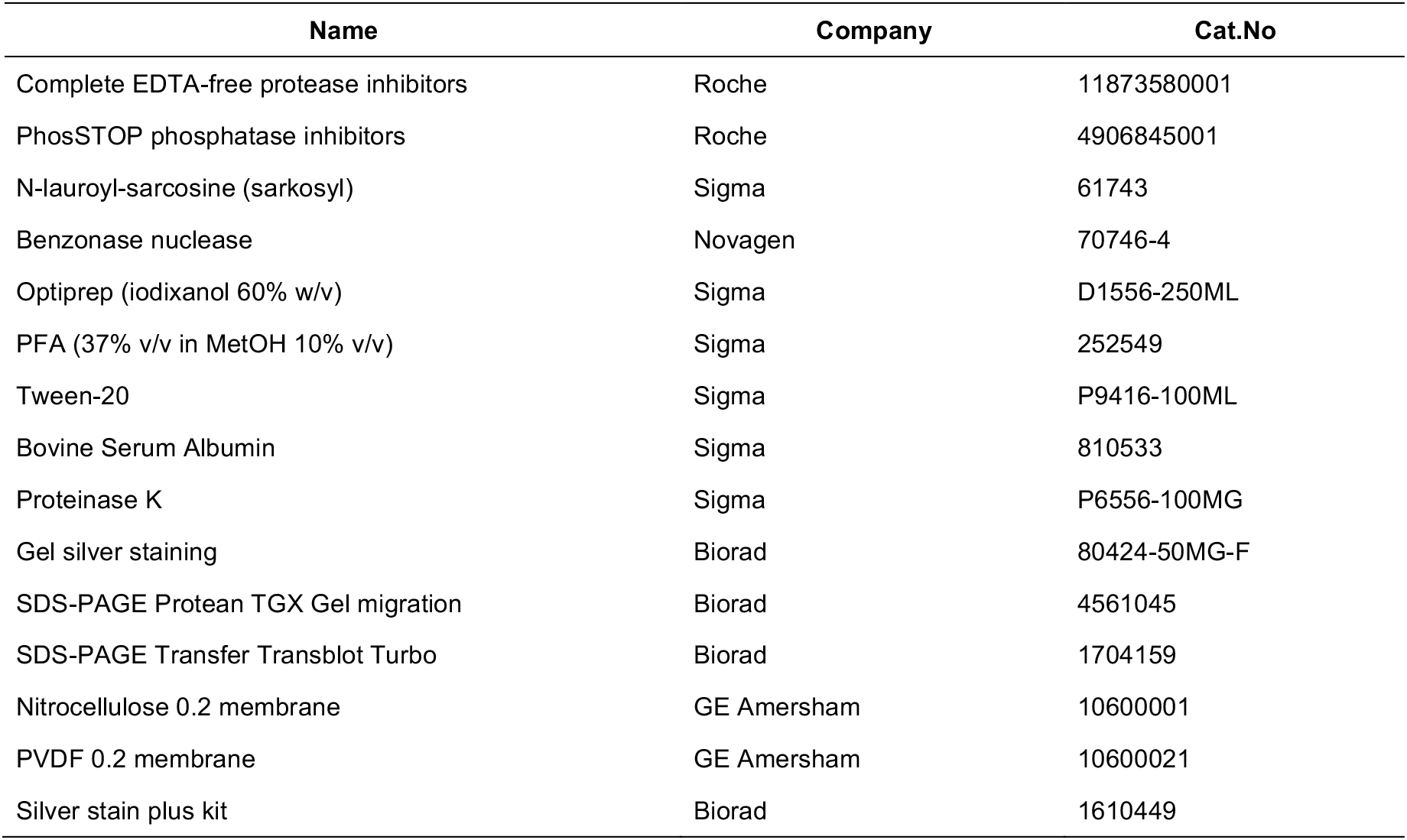

| Name | Company | Cat.No |
| --- | --- | --- |
| Complete EDTA-free protease inhibitors | Roche | 11873580001 |
| PhosSTOP phosphatase inhibitors | Roche | 4906845001 |
| N-lauroyl-sarcosine (sarkosyl) | Sigma | 61743 |
| Benzonase nuclease | Novagen | 70746-4 |
| Optiprep (iodixanol 60% w/v) | Sigma | D1556-250ML |
| PFA (37% v/v in MetOH 10% v/v) | Sigma | 252549 |
| Tween-20 | Sigma | P9416-100ML |
| Bovine Serum Albumin | Sigma | 810533 |
| Proteinase K | Sigma | P6556-100MG |
| Gel silver staining | Biorad | 80424-50MG-F |
| SDS-PAGE Protean TGX Gel migration | Biorad | 4561045 |
| SDS-PAGE Transfer Transblot Turbo | Biorad | 1704159 |
| Nitrocellulose 0.2 membrane | GE Amersham | 10600001 |
| PVDF 0.2 membrane | GE Amersham | 10600021 |
| Silver stain plus kit | Biorad | 1610449 |

## DATA AVAILABILITY

Raw data supporting the results reported in this article are in the figure data files or in the related Appendices and are available upon reasonable request. The entire proteomics dataset has been deposited to the ProteomeXchange Consortium via the PRIDE partner repository with the dataset identifier PXD024998.

## ABBREVIATIONS

α-syn: Alpha-synuclein
CTL: Control
PD: Parkinson’s disease
MSA: Multiple system atrophy
IB: Inclusion body
LB: Lewy body
GCI: Glial cytoplasmic inclusion
MS: Mass spectrometry
GO: Gene ontology
FC: Fold change

## ACKNOWLEDGMENTS

The project was conducted using financial support from the Region Nouvelle Aquitaine, the “Grand Prix” from the Del Duca foundation, and the Innovative Medicines Initiative 2 Joint Undertaking under grant agreement No 116060 (IMPRiND). This Joint Undertaking receives support from the European Union’s Horizon 2020 research and innovation programme and EFPIA. This work is supported by the Swiss State Secretariat for Education, Research and Innovation (SERI) under contract number 17.00038. The opinions expressed and arguments employed herein do not necessarily reflect the official views of these funding bodies.

We are grateful to the NeuroCEB that is run by a consortium of Patients Associations: ARSEP (association for research on multiple sclerosis), CSC (cerebellar ataxias), and France Parkinson for providing human brain samples. We would also like to thank Dr. Benjamin Dehay for sourcing biological material. Finally, ultracentrifugation experiments were performed in the Biochemistry and Biophysics Platform of the Bordeaux Neurocampus at the Bordeaux University, funded by the LABEX BRAIN (ANR-10-LABX-43) with the help of Yann Rufin.

## AUTHOR CONTRIBUTIONS

Conceptualization, F.L. F.D.G. and F.I.; methodology, F.L., F.D.G. and F.I.; investigation, F.L., S.C., F.D.G. and F.I.; biological resources E.B.; writing – original draft preparation, F.L., F.D.G. and F.I.; writing – review and editing, F.L., S.C., E.B., F.D.G. and F.I.; visualization, F.L.; supervision, E.B., F.D.G. and F.I.; funding acquisition, E.B..

## SUPPLEMENTARY FIGURES

**Sup. Fig. 1.**
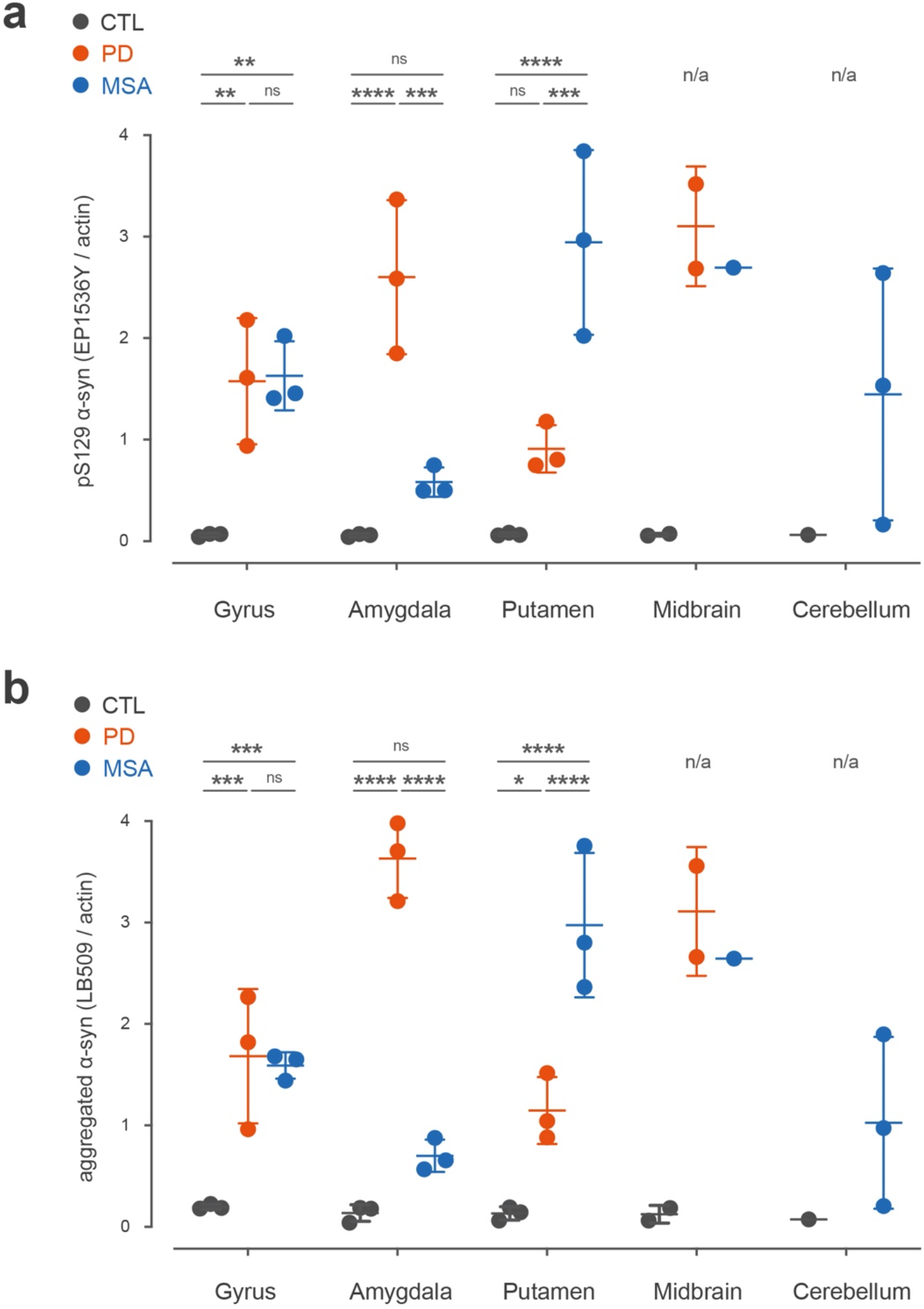
Comparison of pathological alpha-synuclein load in different brain regions of synucleinopathy subjects by immunoblotting. Brain homogenates from different brain regions (gyrus, amygdala, putamen, midbrain and cerebellum) of the three control (black), PD (red) and MSA (blue) human subjects of the study were subjected to dot blot and subsequent immunolabelling and quantification of pS129-positive (**a**, EP1536Y/actin) and aggregated (**b**, LB509/actin) α-synuclein load. The relative amounts (A.U.) are plotted for each subject brain region, and the respective p-values of Tuckey corrected two-way ANOVAs are represented above each couple of comparisons.

**Sup. Fig. 2.**
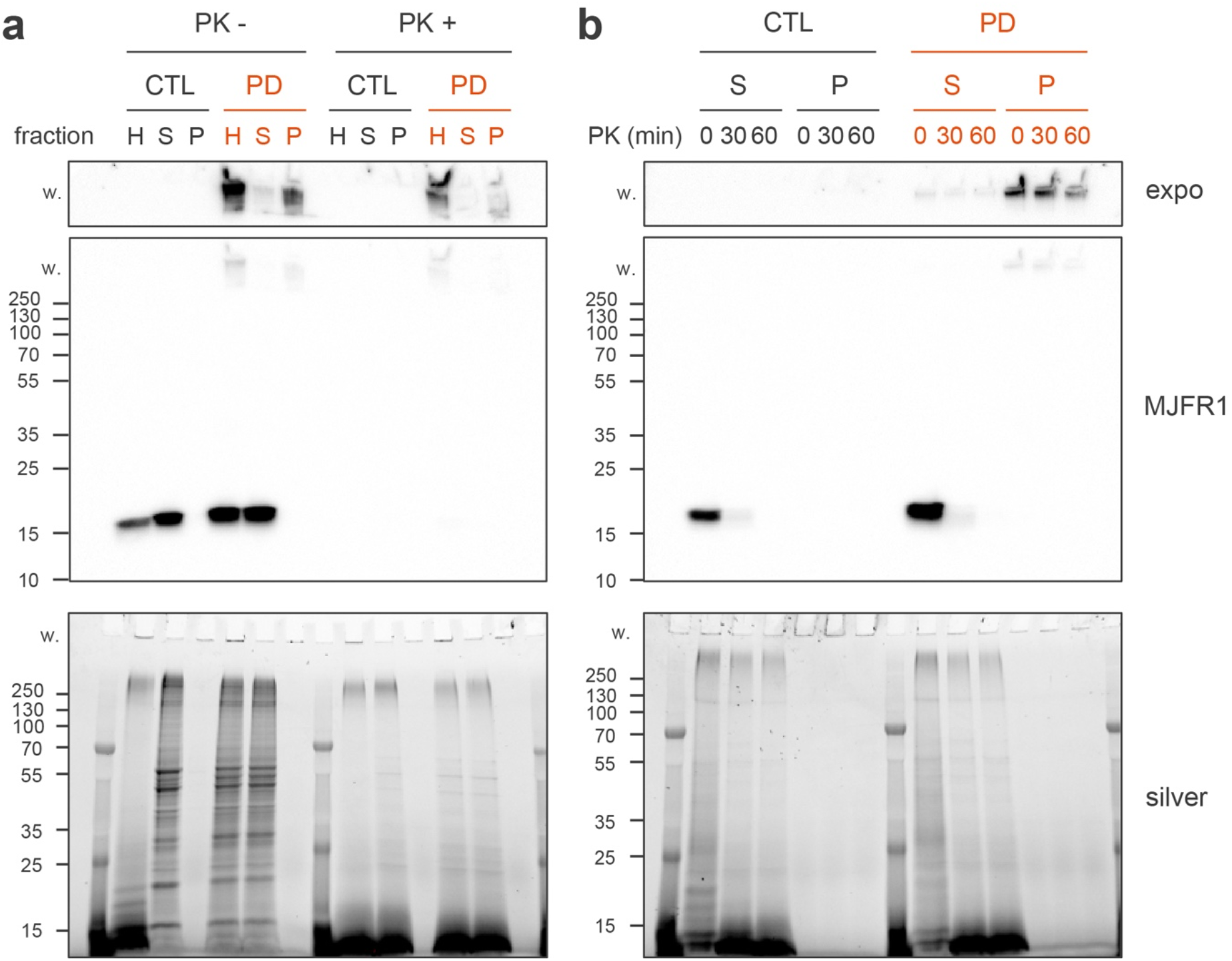
Enrichment in SDS-insoluble, denaturation-PK-resistant high molecular weight alpha-synuclein species in Sarkospin pellets of synucleinopathy brain samples. Biochemical analysis of SDS- and PK-resistance of α-synuclein species contained in Sarkospin fractions extracted from human brain samples. Total solubilized homogenates (H), Sarkospin supernatants (S) and pellets (P) were treated or not for 1 hour (PK+/-, **a**) or treated for the indicated time (0-60 min, **b**) with 1 μg.ml^-1^ PK at 37°C prior to be denatured in Laemmli at 95°C for 5 min, subjected to SDS-PAGE and stained for total proteins (silver) or immunoblotted against human α-synuclein (MJFR1) after transfer on nitrocellulose membrane and fixation. w. indicates the wells and stacking gels were high molecular weight SDS-resistant proteins are retained. Increased signal exposure of these part of membranes are represented in separated boxes (expo.) for better visualization of the latter HMW species.

**Sup. Fig. 3.**
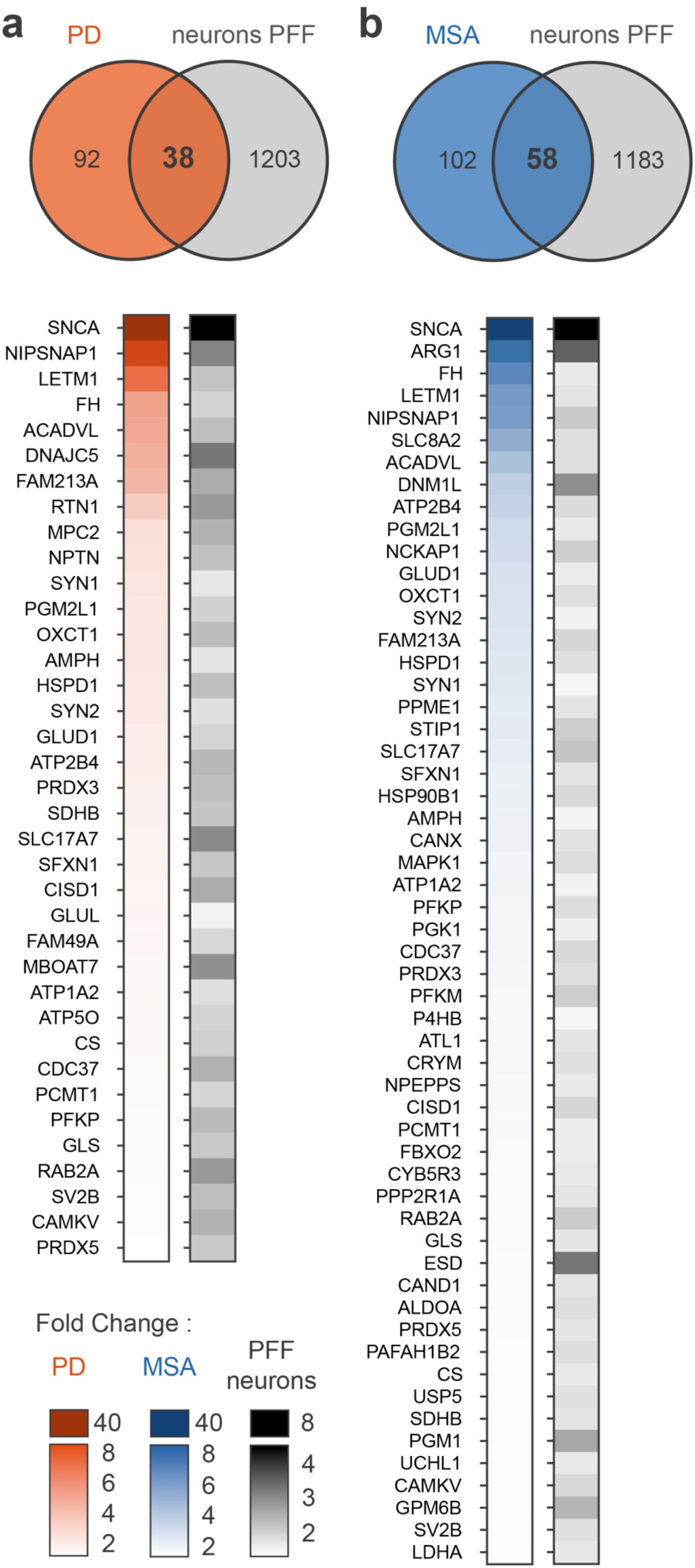
Analysis of the overlaps of insoluble proteomes from synucleinopathy brain samples and α-syn PFF-treated mouse primary neurons^12^. The lists of gene names corresponding to the 130 and 160 proteins enriched in our proteomic study in PD (**a**) and MSA (**b**) pellets, respectively, were searched against the list of gene names of the 1241 unique proteins significantly enriched in insoluble pellets from mouse primary neurons at 14 and 21 dpi treated with recombinant α-syn PFF in Mahul-Mellier et al., 2020^12^. Venn diagrams show that 38 and 58 proteins are common to PD and MSA pellets and PFF-treated neurons insoluble proteins. These proteins are represented in the diagram below, with color-coding their fold change of enrichment PD (red) or MSA (blue) vs CTL, together with their respective mean (14 – 21 dpi) fold change in PFF vs PBS treated mouse primary neurons (grayscale).

**Sup. Fig. 4.**
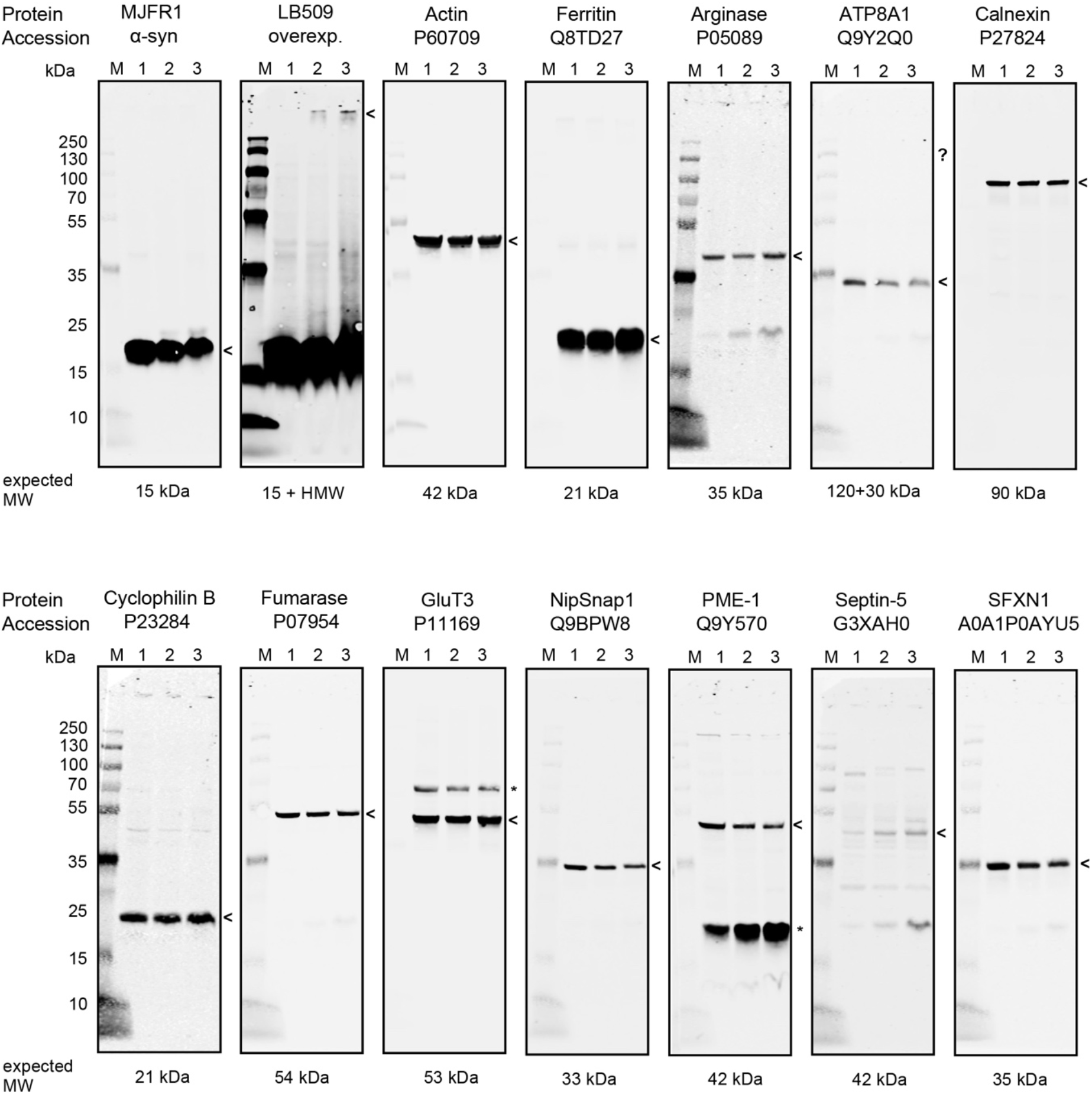
Validation of the selectivity of antibodies directed to candidates of interest by SDS-PAGE. Brain homogenates pools from n=3 controls (lanes 1), PD (lanes 2) and MSA (lanes 3) gyrus samples were denatured by boiling in Laemmli, subjected to SDS-PAGE and blotted on nitrocellulose (lanes M = MW marker). Immunolabelling with antibodies directed to the indicated targets showed bands at the expected size (expected MW, indicated by arrowheads). Stars are pointing at unspecific signals for PME-1, ATP8A1 and GluT3.

**Sup. Fig. 5.**
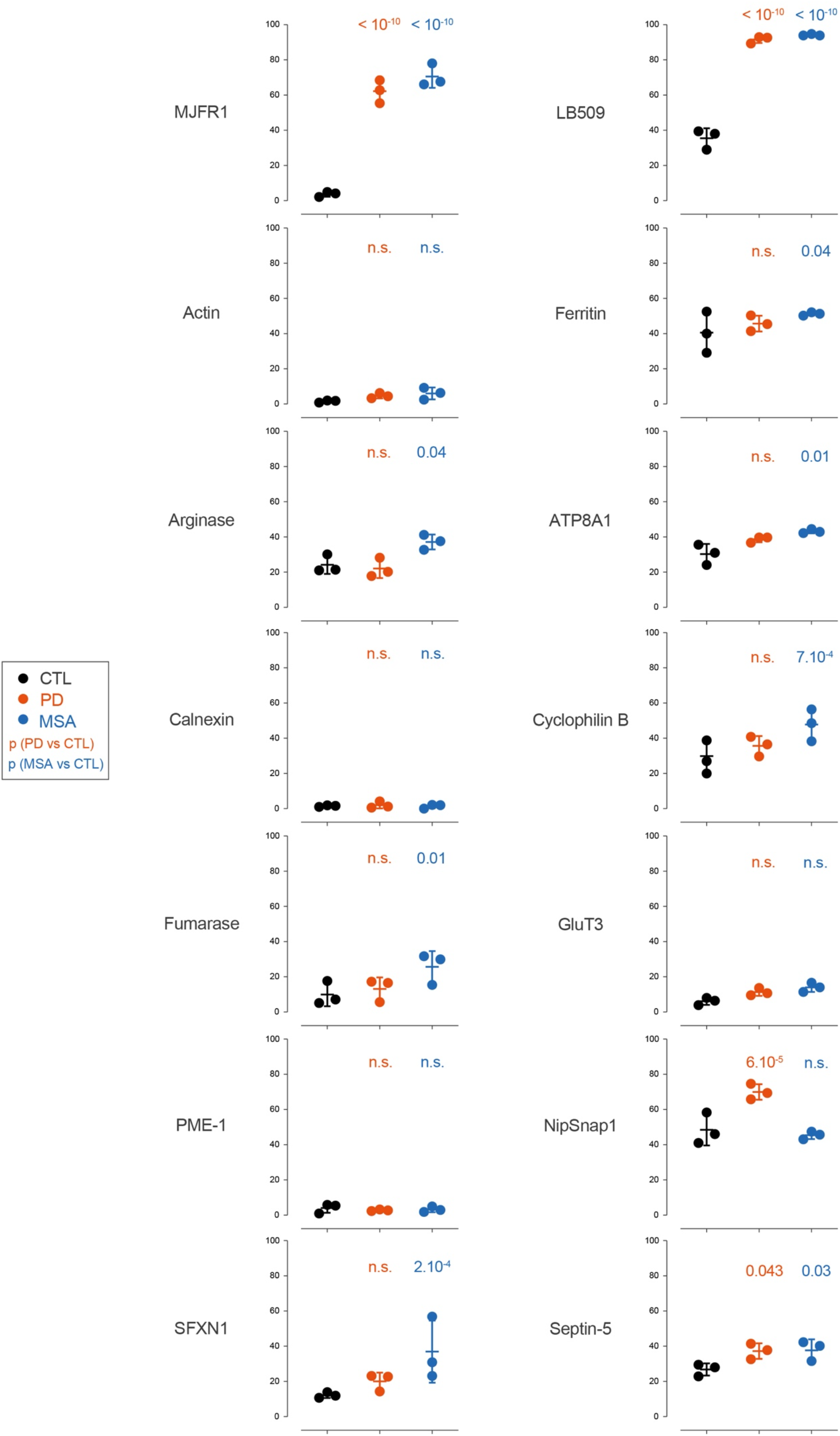
Validation of the enrichment of candidates of interest by dot blot on Sarkospin fractions. Brain homogenates from n=3 controls (black), PD (red) and MSA (blue) gyrus samples were subjected to Sarkospin fractionation. Supernatant and pellet fractions were loaded on drop blot (left) and filter trap (right) and immunolabelled with antibodies directed against the indicated proteins. Each protein amount was quantified in Sarkospin fractions and is plotted as relative pelleted amount (% pellet/total). P-values of the corresponding Tuckey-corrected two-ways ANOVAs of the comparison of insolubility of each target for PD (red) or MSA (blue) vs control, respectively, are shown above the plots. MJFR1 (total α-syn), LB509 (aggregated α-syn), actin and ferritin are shown as controls. PME-1, GluT3 and Calnexin were not validated using these techniques, possibly because of their weak immunodetection, the presence of unspecific bands or isoforms (Sup. Fig. 4), or to the inaccessibility of the epitopes in native conditions of dot blotting.

## APPENDICES

**Appendix A.** List of all proteins detected by mass spectrometry in any Sarkospin pellet.

**Appendix B.** List of 130 proteins enriched in PD.

**Appendix C.** List of 160 proteins enriched in MSA.

**Appendix D.** List of 206 unique proteins gated as enriched in PD and/or MSA.

**Appendix E.** List of 84 gated proteins enriched in both PD and MSA.

**Appendix F.** List of 3 gated proteins specifically enriched only in PD.

**Appendix G.** List of 4 gated proteins specifically enriched only in MSA.

**Appendix H.** List of gene ontology clusters statistically overrepresented in PD and MSA samples.

